# CCR5-del32 is not deleterious in the homozygous state in humans

**DOI:** 10.1101/788117

**Authors:** Daniel Gudbjartsson, Patric Sulem, Kári Stefansson, Nina Mars, Juha Karjalainen, Samuli Ripatti, Aarno Palotie, Mark Daly

## Abstract

Recently, Wei and Nielsen^1^ reported an analysis of UK Biobank data which suggested that the well-known HIV-protective variant CCR5-del32 is associated with a 21% increase in all-cause mortality. We demonstrate, using two well-powered population samples in Iceland and Finland with extensive health data and death information, neither an effect on mortality nor increase in risk of any disease. Further reexamination of the UK Biobank (UKBB) data suggests that the very modest association was with a SNP of poor genotyping quality – at a nearby proxy SNP, no statistically significant impact on mortality nor deviation from Hardy-Weinberg equilibrium exists in the UKBB sample. We thus find no evidence of any meaningful risk of increased mortality from homozygosity of CCR5-del32.

## CCR5-del32 in Iceland

The current freeze of data from the Icelandic project (deCODE genetics) includes 150,828 individuals with both genome wide imputed genotype data and genealogical information. This population sample (53.6% female) had a mean age of 46.7 (SD=18.9) at collection (ongoing since 1997). On average, the individuals have 9.0 years of follow up and 18,259 individuals were deceased as of August 2015. The CCR5 frameshift delta32 variant (hg38 : chr3:46373452: TACAGTCAGTATCAATTCTGGAAGAATTTCCAG : T) is well imputed in the Icelandic set with an imputation information of 0.999. This sequence variant occurs at a similar frequency (minor allele frequency =12.0%) as in Northwestern European populations, resulting in 2,180 homozygotes (genotype frequency 1.44%). This number of homozygotes in Iceland is completely consistent with Hardy Weinberg expectations.

To explore the impact of CCR5-del32 homozygosity on survival, Cox proportional hazard model on survival after blood collection was performed, using sex and age at collection as covariates.

The hazard ratio for death observed for homozygosity was 0.91 (95% CI = 0.80-1.03), if anything slightly protective but not at all significant (P=0.15).

## CCR5-del32 in Finland

The current freeze of data from the FinnGen project includes 135,638 individuals with both genome-wide imputed genotype data and information from a variety of national health registry data including the national death registry. This population sample (56.3% female) currently has a mean age of 60.6 and includes, principally through the inclusion of legacy epidemiology cohorts such as FINRISK and the Finnish Twin Cohort, 11,180 deceased individuals as of December, 2017 (the most recent update of phenotypic information on FinnGen participants). The delta32 variant (hg38 3:46373452:TACAGTCAGTATCAATTCTGGAAGAATTTCCAG:T) is well captured by FinnGen genotyping and imputation with an imputation information score of 0.993 and occurs at a slightly higher frequency (12.9%) than in other European populations, resulting in 2,359 homozygotes – an observed 1.74% of the sample being homozygous in complete consistency with Hardy-Weinberg expectations.

To explore the impact of CCR5-del32 homozygosity on survival, proportional hazards regression was performed on the FinnGen cohort, incorporating sex, 10 principal components of population structure and genotyping batch as covariates, and age on the time scale. The hazard ratio for death observed for homozygosity was 0.95 (0.83-1.09), again, very slightly protective but not significant (p=0.48). Analysis without covariates did not meaningfully change this observation. For technical confirmation, given the nature of the polymorphism, we selected two nearby SNPs a few tens of kb to each side of CCR5-del32 in very high LD (r^2^>0.99 in Finland: rs113010081 and rs113341849) and observed again the same slightly protective trend. Thus the Finnish result, like the Icelandic finding, finds absolutely no evidence of increased mortality from CCR5-del32 homozygosity, in stark contrast to the recently published finding.

## Is there a conflict with the UKBB result?

One might sensibly ask why these results seemingly so clearly conflict with the conclusions of Wei and Nielsen and whether the observations can be reconciled? First and foremost, it should be noted that the original report indicated an effect on increased mortality that was only marginally statistically significant (one-tailed p=.009). This result would of course not stand-up to any reasonable multiple testing correction assuming different endpoints, analytic methods and inheritance models were employed – but regardless, we understand well from the candidate gene era of human genetics that however plausible a hypothesis, any association with this level of statistical evidence of association demands replication before it could be considered true. This is an area where we must maintain even greater vigilance in the modern era in which resources like the UKBB support the analyses of many thousands of disease endpoints and quantitative traits simultaneously. Fortunately, collaboratively minded biobanks, such as those utilized here which have access to broad and longitudinal phenotype data, can be rapidly employed to confirm or refute new hypotheses.

On a technical standpoint, we confirmed using direct analysis of the deletion itself, as well as more distant markers in high LD, lack of any signal of reduced mortality – removing any concern that a technical limitation in the replication studies was responsible for the conflict. As described in detail in a parallel re-analysis of the UKBB data (Maier, Akbari et al, bioRxiv), the basis of the original finding in UKBB was a proxy SNP for CCR5-del32 (rs62625034) which has extremely poor genotype quality – while by contrast the more distant proxy used above showed no significant effect on mortality in the UKBB data.

In summary, we see no evidence of an effect on reduced mortality in CCR5-del32 carriers in two well-powered studies from Iceland and Finland and suggest that the initial report simply constitutes a false positive finding. This is not surprising given the level of significance in the original report which should have been clearly marked and treated as a provisional finding in need of further evaluation. We make no comment about the larger context in which this putative association has been widely cited as a cautionary example – but here make only the point that our shared responsibility as human geneticists is to ensure to the greatest extent possible that only truly durable and incontrovertible association findings are advanced to scientific, medical or public policy discussions – further underscoring the importance of collaboration and consensus in our field.

